# Pharmacologic activation of the G protein-coupled estrogen receptor inhibits pancreatic ductal adenocarcinoma

**DOI:** 10.1101/365668

**Authors:** Christopher A. Natale, Jinyang Li, Tzvete Dentchev, Brian C. Capell, John T. Seykora, Ben Z. Stanger, Todd W. Ridky

## Abstract

Female sex is associated with lower incidence and improved clinical outcomes for many cancer types, including pancreatic ductal adenocarcinoma (PDAC). Although the mechanisms responsible for this sex difference are unknown, recent data suggests nonclassical estrogen signaling through the G Protein-coupled Estrogen Receptor (GPER) is likely involved. Here we used murine syngeneic tumor models and human xenografts to test whether GPER signaling inhibits pancreatic ductal adenocarcinoma (PDAC). Activation of GPER with the specific, small molecule agonist G-1 inhibited PDAC proliferation, depleted c-Myc and programmed death ligand 1 (PD-L1), and increased tumor cell immunogenicity. Systemically delivered G-1 was well tolerated in PDAC bearing mice, significantly prolonged survival, and markedly increased the efficacy of PD-1 targeted immune therapy. We detected GPER protein in a majority of spontaneous human PDAC tumors. These data, coupled with the wide tissue distribution of GPER, and our previous work showing that G-1 inhibits melanoma, suggest that GPER agonists may be useful against many different cancer types.

## Introduction

For many cancers, incidence and age-adjusted mortality rates are lower in females than in males, suggesting that biological differences between the sexes influence tumor initiation, progression, and response to modern therapeutics (1–3). Understanding the mechanisms responsible for these differences may lead to identification of new therapeutic targets for cancer. A growing body of evidence suggests that nonclassical estrogen signaling through the G protein-coupled estrogen receptor (GPER) may be tumor suppressive, including in some cancers that are not traditionally considered sex hormone responsive, such as adrenocortical carcinoma, non-small cell lung cancer, colon carcinoma, osteosarcoma, and cutaneous melanoma (4–9). Consistent with this, we recently showed that systemic administration of a specific small molecule synthetic GPER agonist, named G-1 (10), in mice with therapy-resistant syngeneic melanoma, induced differentiation in tumor cells that inhibited proliferation and rendered tumors more responsive to αPD-1 immune-checkpoint blockade (8). GPER is expressed in many tissues (11) and signaling downstream of GPER is mediated by ubiquitous cellular proteins that mediate cAMP signaling. This led us to consider whether that G-1 may have therapeutic utility as a broadly acting anti-cancer agent effective against GPER expressing cancers.

To test this idea, we turned to pancreatic ductal adenocarcinoma (PDAC), a highly aggressive GPER expressing-cancer that is poorly responsive to current therapy, and a major cause of cancer death in the United States (12). As with many cancers, women have lower PDAC incidence and more favorable outcomes than men, suggesting that estrogen may suppress PDAC (1, 2, 13). Consistent with this, use of estradiol containing oral contraceptives, and history of multiple pregnancies, which correlates with high estrogen exposure, are both associated with decreased PDAC risk (14–16). Further supporting the idea that PDAC is influenced by estrogen are human clinical trials showing that tamoxifen, which is a GPER agonist, extends survival in PDAC patients (17, 18). These data, coupled with lack of clear evidence that nuclear estrogen receptors are expressed and functional in PDAC (19), led us to test whether activation of GPER inhibits PDAC.

Li and colleagues recently generated a library of clonal PDAC tumor cell lines from a genetically engineered mouse model of PDAC, KPCY (‘KPCY’ mice, KRas ^LSL-G12D/+^; Trp53 ^L/+^ or Trp53 ^LSL-R172H/+^; Pdx1-Cre; Rosa-YFP), that faithfully recapitulates the molecular, histological, and clinical features of the human disease (20–22). In syngeneic, immunocompetent C57BL/6 mice, the degree of immune infiltration varies with each cell line. This variability reflects the natural heterogeneity in immune infiltrates observed in human PDAC. Here we used these new models, along with established human PDAC tumor lines, to test whether GPER activation inhibits PDAC, and/or improves PDAC response to immune checkpoint blockade.

## Results and Discussion

To test whether PDAC responds to GPER signaling, we used three genetically defined murine PDAC tumor lines that together represent the heterogeneity in immune infiltration and response to therapy: 6419c5 tumors are associated with minimal CD8+ T cell infiltration and respond poorly to combined cytotoxic and immune therapy, 2838c3 tumors attract robust CD8+ T cell infiltration and respond to therapy, and 6499c4 tumors, which are associated with robust CD8+ T-cell infiltration, but only modest responses to therapy (22).

We first determined that GPER is expressed in all 3 PDAC tumor lines (Figure 1A). Each line then also proved to be highly responsive to G-1. We observed a dose-dependent decrease in proliferation, which was associated with a G_1_-S cell cycle block and corresponding decreases in p-RB, c-Myc. The c-Myc depletion is significant, as c-Myc drives cell proliferation, invasion, and escape from immune surveillance, and is commonly overexpressed in many cancers including PDAC. Consistent with the known role of c-Myc as a positive regulator of the immune checkpoint modulator PD-L1 (23), we also noted that GPER activation depleted PD-L1 (Figure 1B-J), which we predicted would render cells more vulnerable to immune clearance. To test whether these effects were GPER dependent, we treated PDAC cells with a 4-fold molar excess of G-36, a specific GPER antagonist (24), and determined that G-36 blocked the effects of G-1 (Figure 1K). The G-1 induced tumor cell growth arrest was not associated with cell death (Figure 1L).

**Figure 1.**
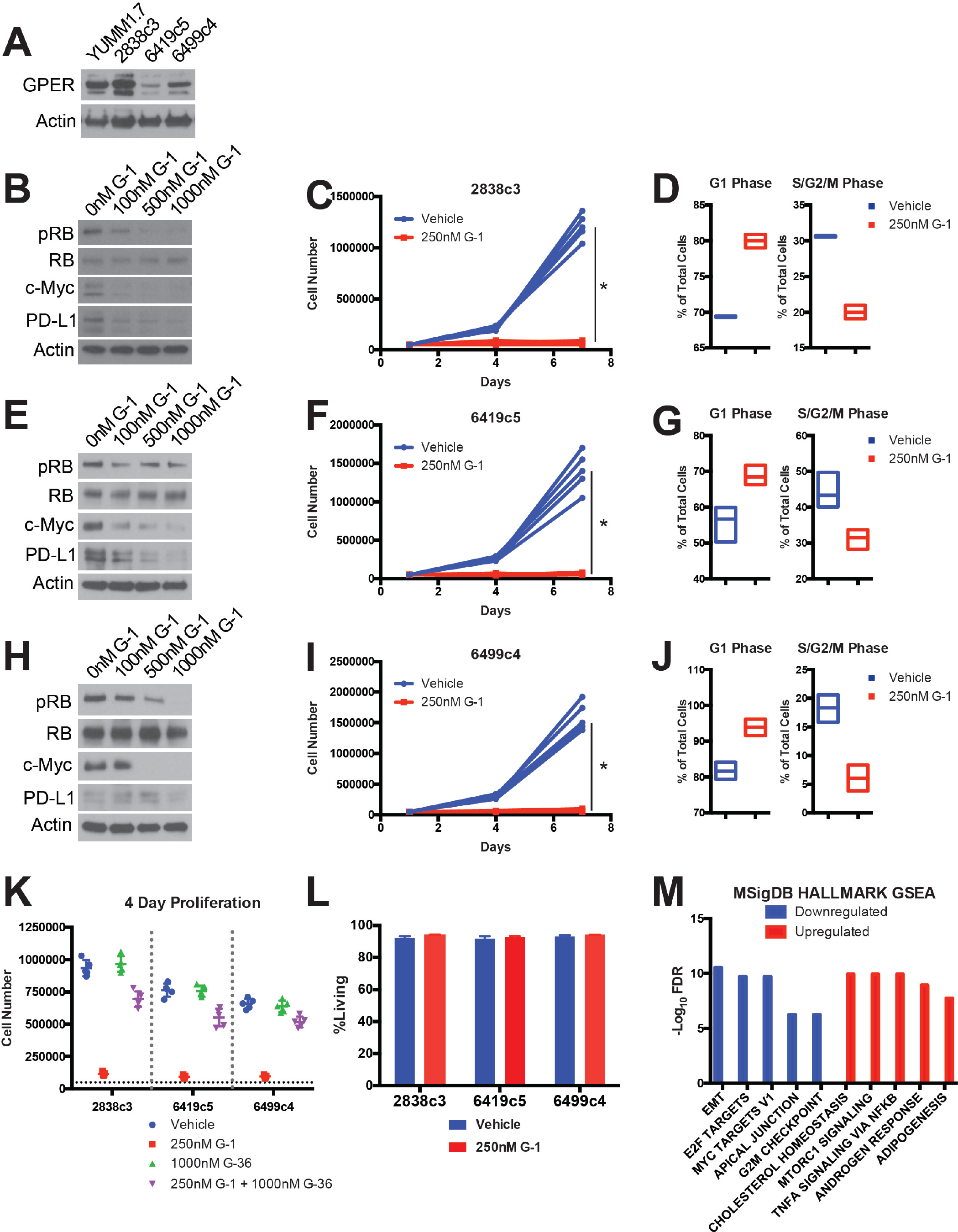
GPER activation inhibits PDAC. (**A**) GPER western blot of lysates from murine YUMM1.7 melanoma cells (GPER positive control) and murine PDAC cells. (**B**) Western analysis of lysates from 2838c3 PDAC cells treated for 16 hours with increasing concentrations of G-1. (**C**) Proliferation of 2838c3 PDAC cells treated with 250nM G-1, n=5 per group, * denotes significance by two-way ANOVA. (**D**) Cell cycle analysis of 2838c3 PDAC cells treated with 250nM G-1, n=3 per group. (**E**) Western analysis of lysates from 6419c5 PDAC cells treated for 16 hours with increasing concentrations of G-1. (**F**) Proliferation of 6419c5 PDAC cells treated with 250nM G-1, n=5 per group, * denotes significance by two-way ANOVA. (**G**) Cell cycle analysis of 6419c5 PDAC cells treated with 250nM G-1, n=3 per group. (**H**) Western analysis of lysates from 6499c4 PDAC cells treated for 16 hours with increasing concentrations of G-1. (**I**) Proliferation of 6499c4 PDAC cells treated with 250nM G-1, n=5 per group, * denotes significance by two-way ANOVA. (**J**) Cell cycle analysis of 6499c5 PDAC cells treated with 250nM G-1, n=3 per group. (**K**) Proliferation assay of 2838c3, 6419c5, and 6499c4 PDAC cells treated with vehicle, 250nM G-1, 1000nM G-36, or a combination of 250nM G-1 and 1000nM G-36. (**L**) Viability assay of murine PDAC cells treated with 250nM G-1, n=3 per group. (**M**) MSigDB HALLMARK gene set enrichment analysis of overlapping upregulated and downregulated genes in 2838c3, 6419c5, and 6499c4 cells treated with 250nM G-1 for 16 hours.

To determine whether GPER induces additional changes in PDAC cells that would suggest general anti-tumor activity, we performed RNA-Seq and HALLMARK gene set enrichment analysis on 2838c3, 6419c5, 6499c4 tumor cells treated with G-1 vs. vehicle control (Figure 1M). Consistent with changes in proliferation and specific proteins in (Figure 1), GPER activation was broadly associated with decreased expression of genes involved in cell proliferation, invasion, and immune evasion including: epithelial-to-mesenchymal transition drivers, E2F targets, c-Myc targets, and cell cycle checkpoint regulators. These data are all consistent with the hypothesis that GPER signaling is tumor suppressive.

We next tested whether G-1 inhibited PDAC in vivo, and whether anti-tumor activity was saturable at pharmacologically achievable, non-toxic doses (Supplementary Figure 1A). We observed tumor responses at 0.1mg/kg (Supplementary Figure 1B), with a maximal response that saturated at 1 mg/kg G-1. We did not observe any deleterious effects at doses up to 100 mg/kg. A pharmacokinetic analysis of at the 10 mg/kg G-1 in mice showed a maximum plasma exposure of 72.4ng/mL (176nM), which is comparable to the saturating exposures we have observed in vitro (Figure 1). We therefore used a dose of 10mg/kg for all subsequent studies, to ensure full GPER activity in vivo.

We next tested whether G-1 might have therapeutic utility as a systemically delivered agent for established PDAC, with and without immune checkpoint inhibitors. Mice harboring syngeneic PDAC were treated with subcutaneously administered G-1, αPD-1 antibody, or both, and tumor growth and survival were compared to matched controls treated with vehicle and isotype antibody controls (Figure 2A-D). All three PDAC tumor models responded to G-1 monotherapy with rapid initial tumor regression and prolonged survival. Tumor response to αPD-1 was different in each tumor line, but was significantly potentiated by G-1 in 2 of the 3 lines. 2838c3 was highly responsive to both G-1 and αPD-1, and the combination of both agents completely cleared tumors in 60% of animals with no evidence of disease at day 100, suggesting a combinatorial benefit. 6419c5 responded to G-1 but was completely resistant to αPD-1 monotherapy. However, G-1 and αPD-1 combination therapy extended survival beyond that observed with G-1 monotherapy, again suggesting combinatorial benefit. In contrast, combination therapy did not provide any additional benefit over G-1 monotherapy in 6499c4 tumors, which were only minimally responsive to αPD-1 alone.

**Figure 2.**
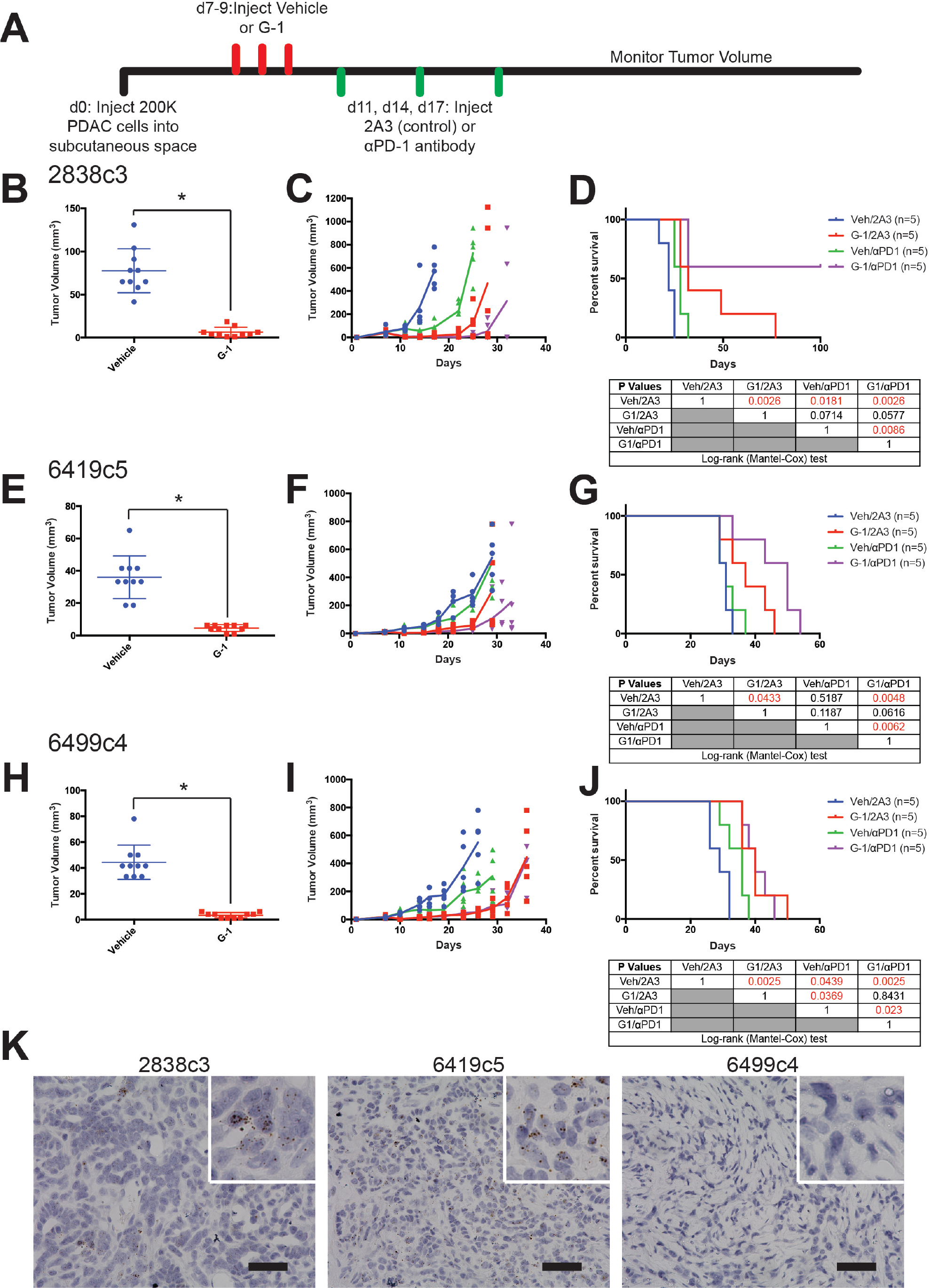
The specific GPER agonist G-1 inhibits murine PDAC in vivo. (**A**) Experimental timeline of murine PDAC-bearing mice treated with subcutaneously delivered vehicle or 10mg/kg G-1, as well as 10mg/kg αPD-1 antibody or isotype antibody control (2A3), n=5 per group. (**B**) Tumor volumes of 2838c3 PDAC tumors one day after the final treatment with 10mg/kg G-1, n=10 per group, * denotes significance by one-way ANOVA. (**C**) 2838c3 tumor volumes measured over time; line terminates after first survival event in the group, n=5 per group. (**D**) Survival curve of 28383-bearing mice treated with vehicle or 10mg/kg G-1, as well as 10mg/kg αPD-1 antibody or 10mg/kg isotype antibody control (2A3), significance between groups by the Log-Rank (Mantel-Cox) test is listed in the table below. (**E**) Tumor volumes of 6419c5 PDAC tumors one day after the final treatment with 10mg/kg G-1, n=10 per group, * denotes significance by one-way ANOVA. (**F**) 6419c5 tumor volumes measured over time; line terminates after first survival event in the group, n=5 per group. (**G**) Survival curve of 6419c5-bearing mice treated with vehicle or 10mg/kg G-1, as well as 10mg/kg αPD-1 antibody or 10mg/kg isotype antibody control (2A3), significance between groups by the Log-Rank (Mantel-Cox) test is listed in the table below. (**H**) Tumor volumes of 6499c4 PDAC tumors one day after the final treatment with 10mg/kg G-1, n=10 per group, * denotes significance by one-way ANOVA. (**I**) 6499c4 tumor volumes measured over time; line terminates after first survival event in the group, n=5 per group. (**J**) Survival curve of 6499c4-bearing mice treated with vehicle or 10mg/kg G-1, as well as 10mg/kg αPD-1 antibody or 10mg/kg isotype antibody control (2A3), significance between groups by the Log-Rank (Mantel-Cox) test is listed in the table below. (**K**) in situ hybridization for PD-L1 in murine PDAC tumors, 40x magnification, scale bar=50um.

In an effort to understand the mechanistic basis for the heterogeneous responses to αPD-1 with or without G-1, we next tested whether PD-L1 is expressed in each tumor. Using *in situ* hybridization for PD-L1 in naïve tumors, we detected high levels of PD-L1 expression in the anti-PD-1 responsive 2838c3 and 6419c5 lines, and complete absence of PD-L1 in the non-responding 6499c4 line (Figure 2E), indicating that the combinatorial survival-promoting effect of G-1 and αPD-1 depends on whether the tumor cells express PD-L1.

Next, we used well-established human PDAC cell lines, harboring the activating K-Ras mutation that drives the vast majority of human PDAC, to test whether GPER activation has similar effects in human models. We detected GPER protein in GPER is expressed in all 3 human lines (Supplementary Figure 2A). To test whether the effects of GPER signaling in human PDAC paralleled those in murine PDAC, we treated the human PDAC tumor cell lines with G-1, and observed similar dose-dependent decreases in p-RB, and c-Myc, which were paralleled by decreases in proliferation (Supplementary Figure 2B-G). We also observed decreased PD-L1 in HPAC and MIA PaCa-2 cells, but did not observe PD-L1 protein in untreated PANC-1 cells. To test whether G-1 has therapeutic utility against human PDAC in vivo, we treated nude mice harboring HPAC and MIA PaCa-2 tumors with subcutaneous G-1 one week after tumor implantation. The PANC-1 cell line failed to establish tumors in nude mice following our protocol. Treatment with G-1 significantly inhibited tumor growth and prolonged survival relative to vehicle treated controls (Supplementary Figure 2H-K). As these studies with human PDAC models were conducted in immunodeficient mice, it was not possible to test for combinatorial activity with immune therapy, as we did in the murine models. Nonetheless, these in vitro and in vivo data using human PDAC are consistent with our findings in mouse models, and together support the idea that GPER activation with G-1 inhibits PDAC.

We next questioned the extent to which GPER is expressed in spontaneous human PDAC. Using a tissue microarray representing several stages of PDAC, immunohistochemical staining for GPER demonstrated both peripheral membrane and punctate cytoplasmic staining, alone or in combination in tumor cells. There was a wide range of staining intensity across different clinical stages (Figure 4A-B). Overall, GPER was detected in 61% of the PDAC cases tested, suggesting that GPER may be a widely expressed, and pharmacologically accessible therapeutic target in human PDAC.

Together with our previous study on melanoma, this work raises the possibility that GPER agonists may have therapeutic utility against a wide array of GPER-expressing cancer types, and critically, may extend the utility of modern immune therapeutics to tumors, such as PDAC, that have thus far been resistant to immune therapy (25). These data highlight the importance of G protein-coupled receptor signaling in cancer, demonstrate that activation of GPER is tumor suppressive in cancers that are not classically hormone responsive, and suggest that GPER activity may contribute to biological differences between the sexes that influence cancer progression and response to modern therapies.

---

**Supplementary Figure 1.**
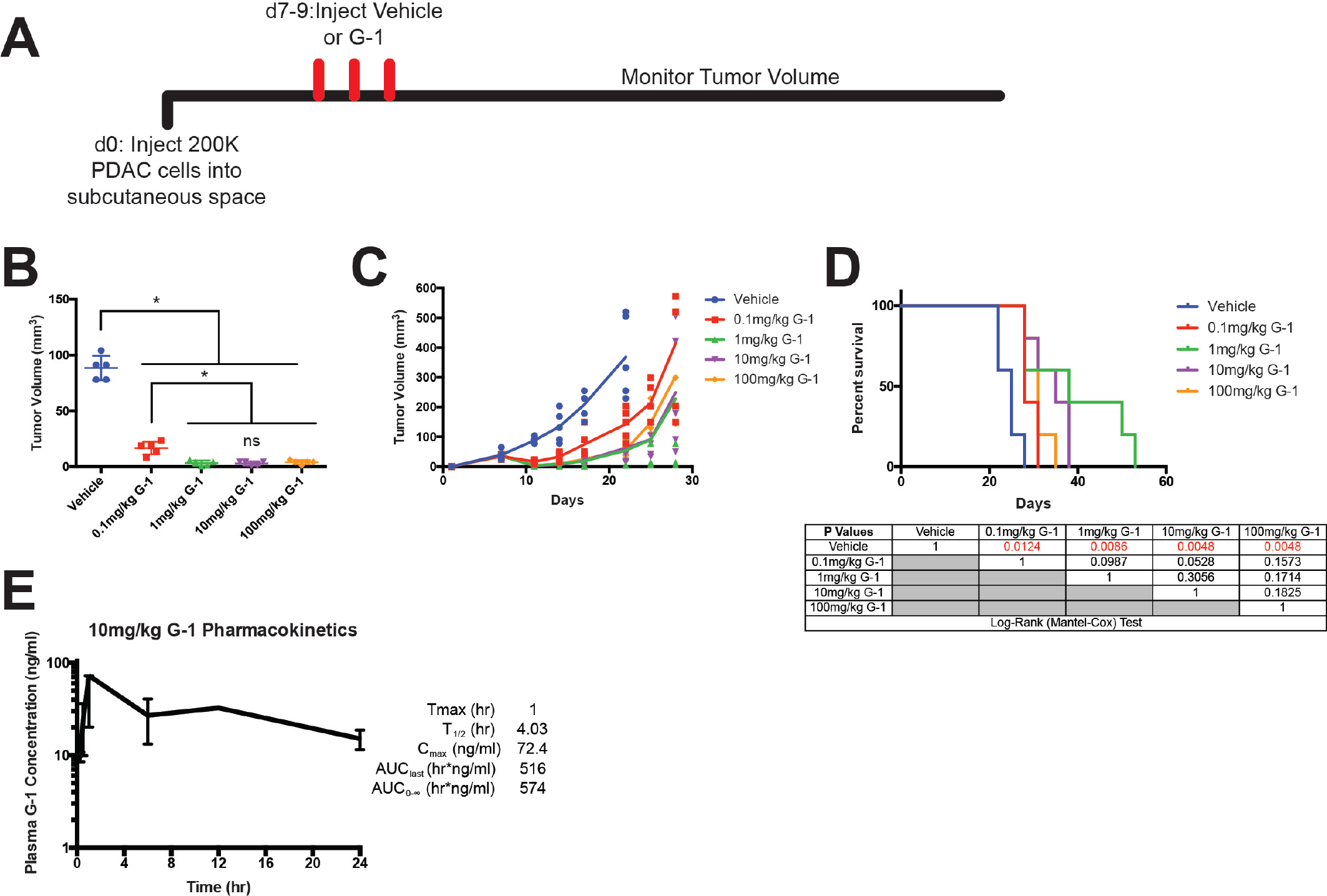
G-1 dose response in vivo and pharmacokinetics. (**A**) Experimental timeline of murine 2838c3 PDAC-bearing mice treated with subcutaneously delivered vehicle or a dose response of G-1, n=5 per group. (**B**) Tumor volumes of 2838c3 PDAC tumors one day after the final treatment with vehicle or G-1, n=5 per group, * denotes significance by one-way ANOVA. (**C**) 2838c3 tumor volumes measured over time; line terminates after first survival event in the group, n=5 per group. (**D**) Survival curve of 28383-bearing mice treated with vehicle or a dose response of G-1, significance between groups by the Log-Rank (Mantel-Cox) test is listed in the table below. (**E**) Pharmacokinetics of 10mg/kg G-1 in mice, n=3 mice per time point.

**Supplementary Figure 2.**
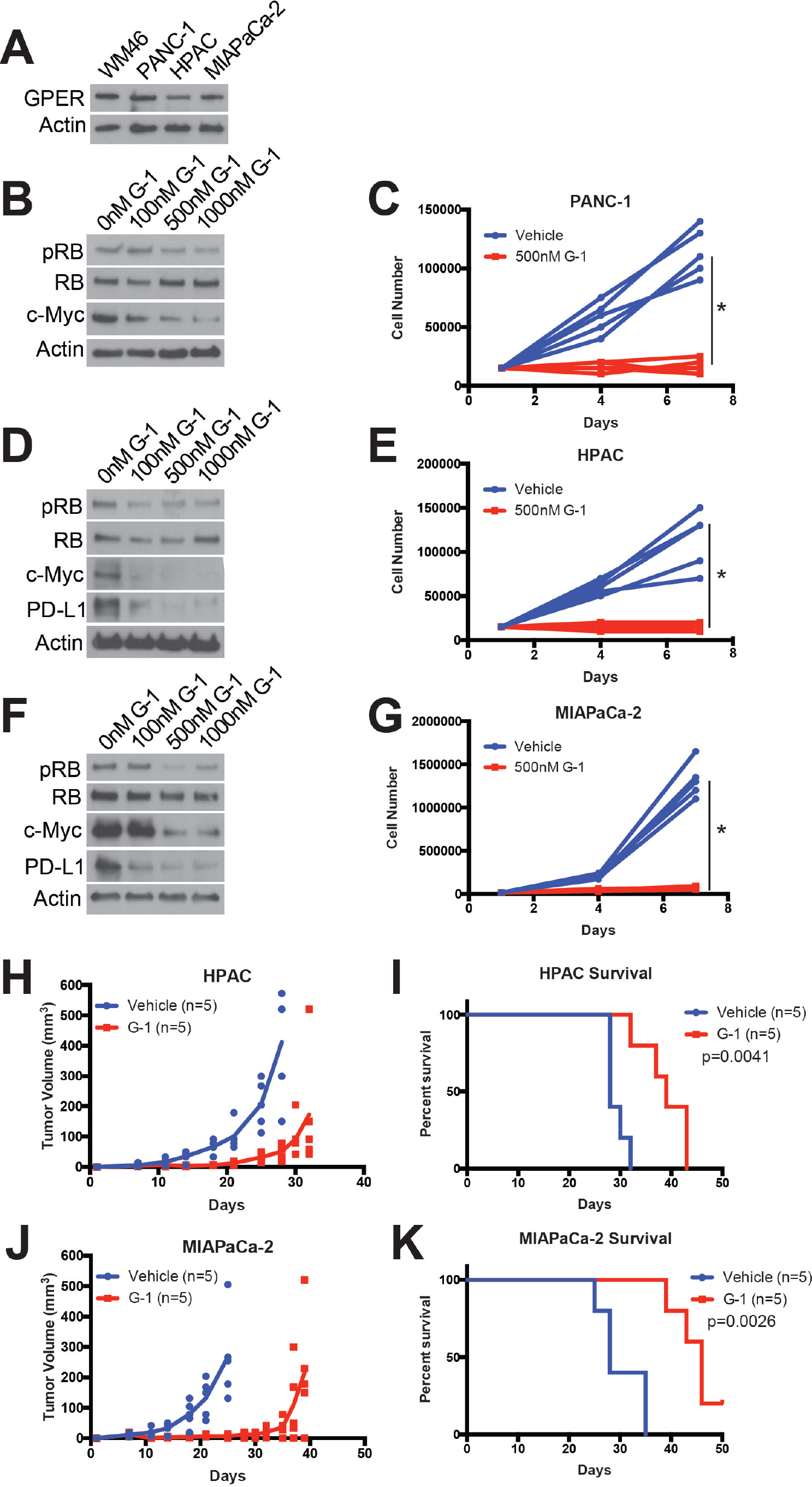
The specific GPER agonist G-1 inhibits human PDAC. (**A**) GPER western blot of lysates from human WM46 melanoma cells (GPER positive control) and human PDAC cells. (**B**) Western analysis of lysates from PANC-1 PDAC cells treated for 16 hours with increasing concentrations of G-1. (**C**) Proliferation of PANC1 PDAC cells treated with 500nM G-1, n=5 per group, * denotes significance by two-way ANOVA. (**D**) Western analysis of lysates from HPAC PDAC cells treated for 16 hours with increasing concentrations of G-1. (**E**) Proliferation of HPAC PDAC cells treated with 500nM G-1, n=5 per group, * denotes significance by two-way ANOVA. (**F**) Western analysis of lysates from MIA PaCa-2 PDAC cells treated for 16 hours with increasing concentrations of G-1. (**G**) Proliferation of MIA PaCa-2 PDAC cells treated with 500nM G-1, n=5 per group, * denotes significance by two-way ANOVA. (**H**) HPAC tumor volumes measured over time; line terminates after first survival event in the group, n=5 per group. (**I**) Survival curve of HPAC-bearing mice treated with vehicle or 10mg/kg G-1 on day 7-9, 14-16, and 21-23 (3 weekly pulses), significance between groups by the Log-Rank (Mantel-Cox) test. (**J**) MIA PaCa-2 tumor volumes measured over time; line terminates after first survival event in the group, n=5 per group. (**K**) Survival curve of MIA PaCa-2-bearing mice treated with vehicle or 10mg/kg G-1 on day 7-9, 14-16, and 21-23 (3 weekly pulses), significance between groups by the Log-Rank (Mantel-Cox) test.

**Supplementary Figure 3.**
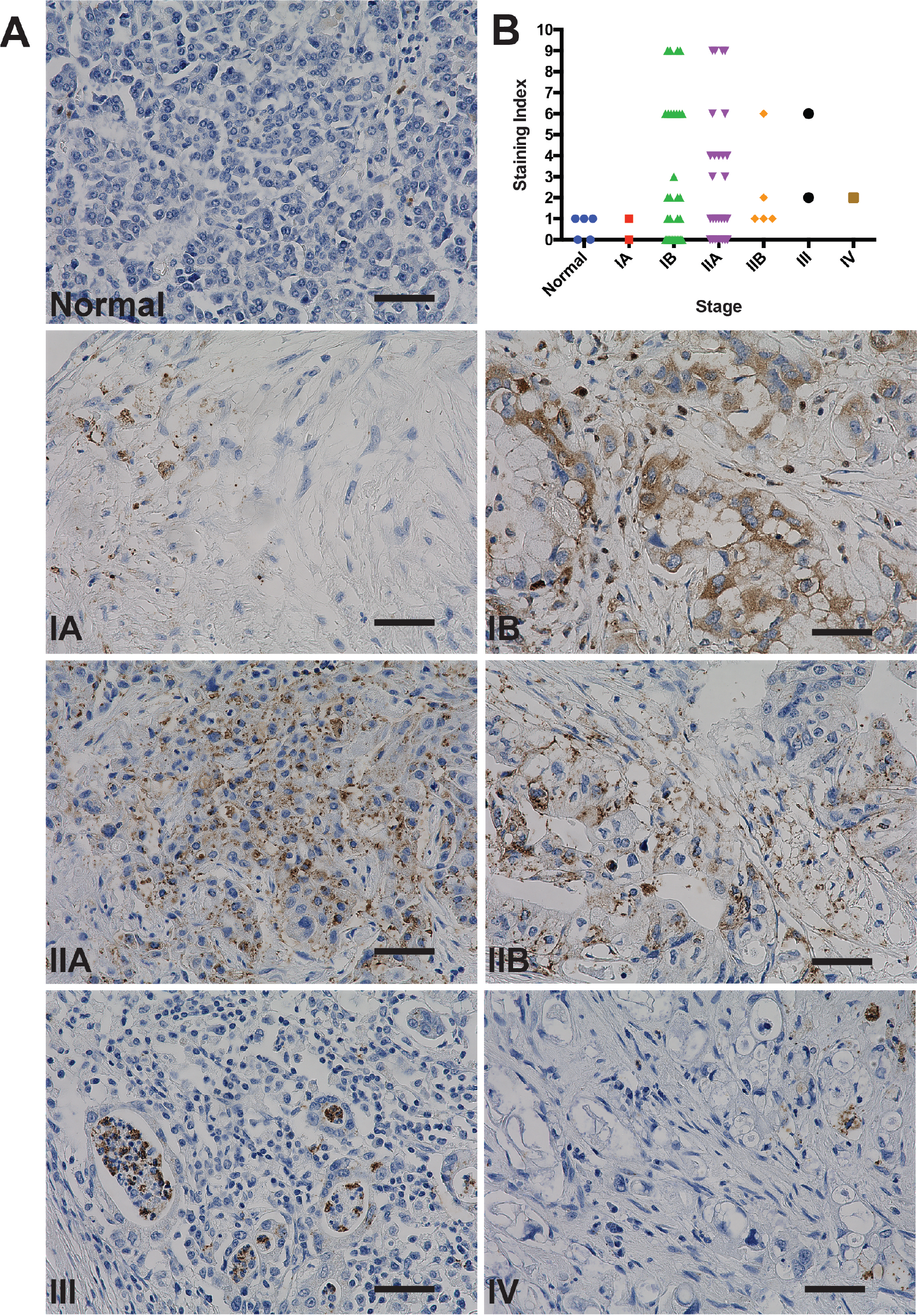
GPER is expressed in human PDAC clinical samples. (**A**) Representative images of normal human pancreas and stage 1A-IV PDAC stained for GPER, 40x magnification, scale bar=50um. (**B**) Pathologist scoring of the pancreatic tissue microarray stained for GPER, scoring index was determined by scoring the percentage of positive cells on a scale of 0 to 3, as well as the intensity of GPER staining on a scale of 0 to 3, these scores were multiplied to generate the scoring index.

## Methods

### Cell culture and cell lines

2838c3, 6419c5, 6499c4 murine PDAC cells were derived in the laboratory of Ben Stanger (University of Pennsylvania) and cultured in DMEM (Mediatech, Manassas, VA, USA) with 5% FBS (Invitrogen, Carlsbad, CA, USA) and 1% antibiotic-antimycotic (Invitrogen). PANC-1, HPAC, and MIA PaCa-2 cell lines were a gift from the laboratory of Ben Stanger (University of Pennsylvania) and cultured in DMEM (Mediatech) with 5% FBS (Invitrogen) and 1% antibiotic-antimycotic (Invitrogen). WM46 melanoma cells were a gift from Meenhard Herlyn (Wistar Institute, Philadelphia, PA, USA) and were cultured in TU2% media. Tumor cells were regularly tested using MycoAlert Mycoplasma Detection Kit from Lonza (Allendale, NJ, USA). G-1 (10008933) and G-36 (14397) were purchased from Cayman Chemical (Ann Arbor, MI, USA). Cells were trypsinized using 0.05% Trypsin with EDTA (Invitrogen) for 5 minutes to detach from the plate.

### Mice

All mice were purchased from Jackson Laboratories (Bar Harbor, ME, USA). Five to seven week old female C57BL/6J or nude (NU/J) mice were allowed to acclimatize for one week prior to being used for experiments. These studies were preformed without inclusion/exclusion criteria or blinding, but included randomization. Based on a twofold-anticipated effect, we performed experiments with at least 5 biological replicates. All procedures were performed in accordance with International Animal Care and Use Committee (IACUC)-approved protocols at the University of Pennsylvania.

### Subcutaneous tumors and treatments

Subcutaneous tumors were initiated by injecting tumor cells in 50% Matrigel (Corning, Bedford, MA, USA) into the subcutaneous space on the left and right flanks of mice. 2 × 10^5^ of murine PDAC cells or 5 × 10^5^ human PDAC cells were used for each tumor. In vivo G-1 treatments were performed by first dissolving G-1, synthesized as described previously (9), in 100% ethanol at a concentration of 1mg/ml. The desired amount of G-1 was then mixed with an appropriate volume of sesame oil, and the ethanol was evaporated off using a Savant Speed Vac (Thermo Fisher Scientific, Waltham, MA, USA), leaving the desired amount of G-1 dissolved in 50µL of sesame oil per injection at a 10mg/kg dose. Vehicle injections were prepared in an identical manner using 100% ethanol. Vehicle and G-1 injections were delivered through subcutaneous injection as indicated in each experimental timeline. Isotype control antibody (Clone: 2A3, BioXcell, West Lebanon, NH, USA) and αPD-1 antibody (Clone: RMP1-14, BioXcell) were diluted in sterile PBS and delivered through intraperitoneal injections at a dose of 10mg/kg.

### Survival Analysis

As subcutaneous tumors grew in mice, perpendicular tumor diameters were measured using calipers. Volume was calculated using the formula L × W^2 × 0.52, where L is the longest dimension and W is the perpendicular dimension. Animals were euthanized when tumors exceeded a protocol-specified size of 15 mm in the longest dimension. Secondary endpoints include severe ulceration, death, and any other condition that falls within the IACUC guidelines for Rodent Tumor and Cancer Models at the University of Pennsylvania.

### Western Blot Analysis

Adherent cells were washed once with DPBS, and lysed with 8M urea containing 50mM NaCl and 50mM Tris-HCl, pH 8.3, 10mM dithiothreitol, 50mM iodoacetamide. Lysates were quantified (Bradford assay), normalized, reduced, and resolved by SDS gel electrophoresis on 4–15% Tris/Glycine gels (Bio-Rad, Hercules, CA, USA). Resolved protein was transferred to PVDF membranes (Millipore, Billerica, MA, USA) using a Semi-Dry Transfer Cell (Bio-Rad), blocked in 5% BSA in TBS-T and probed with primary antibodies recognizing β-Actin (Cell Signaling Technology, #3700, 1:4000, Danvers, MA, USA), c-Myc (Cell Signaling Technology, #13987, 1:1000), GPER (Sigma, HPA027052, 1:500), HLA-ABC (Biolegend, w6/32,1:500, San Diego, CA, USA), human PD-L1 (Cell Signaling Technology, #13684, 1:1000), mouse PD-L1 (R&D systems, AF1019, 1:500, Minneapolis, MN, USA), p-RB (Cell Signaling Technology, #8516, 1:1000), and RB (Cell Signaling Technology, #9313, 1:1000). After incubation with the appropriate secondary antibody, proteins were detected using either Luminata Crescendo Western HRP Substrate (Millipore) or ECL Western Blotting Analysis System (GE Healthcare, Bensalem, PA). All western blots were repeated at least 3 times.

### Immunohistochemistry and Quantification

FFPE tissue microarrays were purchased from Biomax (9461e, Derwood, MD, USA) and were stained GPER (Novus Biologics, NLS1183, Littleton, CO, USA) as previously described with some modifications (8). Briefly, slides have been deparaffinized and rehydrated in extend time than standard immunohistochemistry protocol (in three xylenes 7 min each time, 3 times 100% alcohol, 2 time 95% alcohol, ones 70% and 50% alcohol and finished with distill water). The antigen retrieval was done by immersing the slides in Tris-EDTA pH 8.0 and microwave for 14 minutes at power 9 then cool down to room temperature on the bench, washed three times in wash buffer and blocked sequentially the endogenous peroxidase and non-specific protein binding. The volume and the dilution/concentration (500 ul per slide at the dilution 1:70) of the primary antibody was calculate considering the total area of tissue and incubate overnight at 40C. Following multiple washes secondary antibody, goat anti-rabbit conjugated to HRP was applied, incubated for 20 minutes at room temperature, then washed out and signal was amplified with substrate-DAB chromogen buffer. The tissues were counterstain with hematoxylin, dehydrated, cover-slipped and analyzed. A board-certified pathologist performed scoring of the stained tissue microarray, and scoring index was determined by scoring the percentage of positive cells on a scale of 0 to 3 as well as the intensity of GPER staining on a scale of 0 to 3, these scores were multiplied to generate the scoring index value.

### Cell cycle analysis

Tumor cells were cultured in 5% FBS in DMEM following standard cell culture protocol. Hoechst 33342 (ThermoFisher Scientific, #H1399) was added directly to the cell culture medium with the final concentration of 10 ug/mL 1 hour before sample collection. Cells were incubated with Hoechst 33342 at 37C for 1 hour. Then, cells were prepared to single cell suspension using trypsin and washed with PBS twice. Cells were resuspended in flow buffer (1% FBS in PBS) for flow analysis in a BD LSR II flow cytometry machine. Flow results were analyzed using the FlowJo software to assess the percentage of cells in G1 phase and S-G2-M phase.

### Cell viability analysis

Tumor cells were cultured in 5% FBS in DMEM following standard cell culture protocol. Cells were prepared to single cell suspension using trypsin and washed with PBS twice. Cells were resuspended in flow buffer (1% FBS in PBS) with DAPI (ThermoFisher Scientific, #D21490) incubation for flow analysis in a BD LSR II flow cytometry machine. Flow results were analyzed using the FlowJo software to assess the percentage of cells with negative staining of DAPI.

### RNA-seq

RNA was extracted using RNeasy kit (Qiagen catalog no. 74014) following the manufacturer’s instructions. All RNA-seq libraries were prepared using the NEBNext Poly(A) mRNA Magnetic Isolation Module followed by NEBNext Ultra Directional RNA Library Prep Kit for Illumina (both from NEB). Library quality was analyzed using Agilent BioAnalyzer 2100 (Agilent) and libraries were quantified using NEB Library Quantification Kits (NEB). Libraries were then sequenced using a NextSeq500 platform (75bp, single-end reads) (Illumina). All RNA-seq was aligned using RNA STAR under default settings to Homo sapiens UCSC hg19 (RefSeq & Gencode gene annotations). FPKM generation and differential expression analysis was performed using DESeq2. DESeq2 output was analyzed by comparing differentially expressed genes with p<0.05 and HALLMARK gene set enrichment analysis was performed using MSigDB database (26).

### RNA in situ hybridization analysis

Tumor cells were subcutaneously implanted into C57BL/6 mice for tumor growth. Tumors were collected 3 weeks after implantation, fixed with Zinc Formalin Fixative (Polysciences, #21516), and embedded in paraffin. Tumor sample paraffin sections were used for RNA in situ hybridization analysis using the RNAscope 2.5 HD Assay – BROWN (Advanced Cell Diagnosis) with probe targeting CD274 (PD-L1) (420501). The RNA in situ hybridization analysis was performed following standard procedures from the kit manual and published protocol (27) (RNAscope: a novel in situ RNA analysis platform for formalin-fixed, paraffin-embedded tissues).

### Pharmacokinetic Analysis

Pharmacokinetic Analysis was performed by Ricerca Biosciences (Concord, OH, USA). Briefly, animals were not fasted prior to dosing. Animals were divided into 6 subgroups of 3 animals each. Each subgroup was bled at one time point and terminated following blood collection. G-1 was administered as described in “*Subcutaneous tumors and treatments”* Blood samples were collected from the retrorbital sinus at 0.25, 0.5, 1, 6, 12, and 24 hours post dose. No animal was found dead or deemed moribund during the study. No abnormalities were observed at the detailed clinical observations, during the daily observations, or following initiation of dosing. At each blood collection period, one subgroup of animals was placed under deep anesthesia induced with CO2/O2; while still under anesthesia, the final blood sample was collected and the animals were terminated by cervical dislocation. Plasma samples were sent to Ricerca Bioanalytical department for analysis. Pharmacokinetic analysis was conducted using WinNonlin Version 6.2 (Pharsight, Mountain View, CA, USA), operating as a validated software system. Noncompartmental analysis was conducted using an extravascular administration model. The area under the plasma concentration-time curves, peak plasma concentration, the time to achieve peak plasma concentration, and the plasma terminal half-life (AUC, Cmax, Tmax, and T1/2) were calculated from mean plasma concentrations for each sampling time for G-1. Nominal blood collection times were used for toxicokinetic calculations.

### Statistical Analysis

All statistical analysis was performed using Graphpad Prism 8 (Graphpad Software, La Jolla, CA, USA). No statistical methods were used to predetermine sample size. Details of each statistical test used are included in the figure legends.

## Acknowledgements

The authors thank the University of Pennsylvania Skin Biology and Disease Research-based center for analysis of tissue sections. The authors also thank Michael Feigin, Miriam Doepner, and Junqian Zhang for critical pre-submission review. T.W.R. is supported by a grant from the NIH/NCI (R01 CA163566), a Penn/Wistar Institute N.I.H. SPORE (P50CA174523), the Melanoma Research Foundation, and the Dermatology Foundation. C.A.N was supported by an NIH/NIAMS training grant (T32 AR0007465-32) and an NIH/NCI F31 NRSA Individual Fellowship (F31 CA206325). This work was supported in part by the Penn Skin Biology and Diseases Resource-based Center (P30-AR069589). This work was also supported in part by a phase I STTR from the NIH/NCI (R41CA228695). The contents are solely the responsibility of the authors and do not necessarily represent the official views of the NIH.

## Ethics Statement

This study was performed in strict accordance with the recommendations in the Guide for the Care and Use of Laboratory Animals of the National Institutes of Health. All of the animals were handled according to approved institutional animal care and use committee (IACUC) protocol (#803381) of the University of Pennsylvania.

## Competing Interests

C.A.N. is listed as an inventor on a patent application held by the University of Pennsylvania related to this work (PCT/US2017/035278) and is a cofounder and current employee of the Penn Center for Innovation supported startup Linnaeus Therapeutics Inc. T.W.R. is listed as an inventor on a patent application (PCT/US2017/035278) held by the University of Pennsylvania related to this work, and is a cofounder and consultant to the Penn Center for Innovation supported startup Linnaeus Therapeutics Inc.

The authors have no additional financial interests.

## References

1. Cook MB, Dawsey SM, Freedman ND, Inskip PD, Wichner SM, Quraishi SM, Devesa SS, McGlynn KA. Sex disparities in cancer incidence by period and age. Cancer Epidemiol Biomarkers Prev. 2009;18(4):1174–82. doi: 10.1158/1055-9965.EPI-08-1118. PubMed PMID: 19293308; PMCID: PMC2793271.

2. Cook MB, McGlynn KA, Devesa SS, Freedman ND, Anderson WF. Sex disparities in cancer mortality and survival. Cancer Epidemiol Biomarkers Prev. 2011;20(8):1629–37. doi: 10.1158/1055-9965.EPI-11-0246. PubMed PMID: 21750167; PMCID: PMC3153584.

3. McQuade JL, Daniel CR, Hess KR, Mak C, Wang DY, Rai RR, Park JJ, Haydu LE, Spencer C, Wongchenko M, Lane S, Lee DY, Kaper M, McKean M, Beckermann KE, Rubinstein SM, Rooney I, Musib L, Budha N, Hsu J, Nowicki TS, Avila A, Haas T, Puligandla M, Lee S, Fang S, Wargo JA, Gershenwald JE, Lee JE, Hwu P, Chapman PB, Sosman JA, Schadendorf D, Grob JJ, Flaherty KT, Walker D, Yan Y, McKenna E, Legos JJ, Carlino MS, Ribas A, Kirkwood JM, Long GV, Johnson DB, Menzies AM, Davies MA. Association of body-mass index and outcomes in patients with metastatic melanoma treated with targeted therapy, immunotherapy, or chemotherapy: a retrospective, multicohort analysis. Lancet Oncol. 2018. doi: 10.1016/S1470-2045(18)30078-0. PubMed PMID: 29449192.

4. Zhu G, Huang Y, Wu C, Wei D, Shi Y. Activation of G-Protein-Coupled Estrogen Receptor Inhibits the Migration of Human Nonsmall Cell Lung Cancer Cells via IKK-beta/NF-kappaB Signals. DNA Cell Biol. 2016;35(8):434–42. doi: 10.1089/dna.2016.3235. PubMed PMID: 27082459.

5. Liu Q, Chen Z, Jiang G, Zhou Y, Yang X, Huang H, Liu H, Du J, Wang H. Epigenetic down regulation of G protein-coupled estrogen receptor (GPER) functions as a tumor suppressor in colorectal cancer. Mol Cancer. 2017;16(1):87. doi: 10.1186/s12943-017-0654-3. PubMed PMID: 28476123; PMCID: PMC5418684.

6. Chimento A, Sirianni R, Casaburi I, Zolea F, Rizza P, Avena P, Malivindi R, De Luca A, Campana C, Martire E, Domanico F, Fallo F, Carpinelli G, Cerquetti L, Amendola D, Stigliano A, Pezzi V. GPER agonist G-1 decreases adrenocortical carcinoma (ACC) cell growth in vitro and in vivo. Oncotarget. 2015;6(22):19190–203. doi: 10.18632/oncotarget.4241. PubMed PMID: 26131713; PMCID: PMC4662484.

7. Wang Z, Chen X, Zhao Y, Jin Y, Zheng J. G-protein-coupled estrogen receptor suppresses the migration of osteosarcoma cells via post-translational regulation of Snail. J Cancer Res Clin Oncol. 2018. doi: 10.1007/s00432-018-2768-4. PubMed PMID: 30341688.

8. Natale CA, Li J, Zhang J, Dahal A, Dentchev T, Stanger BZ, Ridky TW. Activation of G protein-coupled estrogen receptor signaling inhibits melanoma and improves response to immune checkpoint blockade. Elife. 2018;7. doi: 10.7554/eLife.31770. PubMed PMID: 29336307; PMCID: PMC5770157.

9. Natale CA, Duperret EK, Zhang J, Sadeghi R, Dahal A, O’Brien KT, Cookson R, Winkler JD, Ridky TW. Sex steroids regulate skin pigmentation through nonclassical membrane-bound receptors. Elife. 2016;5. doi: 10.7554/eLife.15104. PubMed PMID: 27115344; PMCID: PMC4863824.

10. Bologa CG, Revankar CM, Young SM, Edwards BS, Arterburn JB, Kiselyov AS, Parker MA, Tkachenko SE, Savchuck NP, Sklar LA, Oprea TI, Prossnitz ER. Virtual and biomolecular screening converge on a selective agonist for GPR30. Nature chemical biology. 2006;2(4):207–12. doi: 10.1038/nchembio775. PubMed PMID: 16520733.

11. Olde B, Leeb-Lundberg LM. GPR30/GPER1: searching for a role in estrogen physiology. Trends Endocrinol Metab. 2009;20(8):409–16. doi: 10.1016/j.tem.2009.04.006. PubMed PMID: 19734054.

12. Rahib L, Smith BD, Aizenberg R, Rosenzweig AB, Fleshman JM, Matrisian LM. Projecting cancer incidence and deaths to 2030: the unexpected burden of thyroid, liver, and pancreas cancers in the United States. Cancer research. 2014;74(11):2913–21. doi: 10.1158/0008-5472.CAN-14-0155. PubMed PMID: 24840647.

13. Ma J, Siegel R, Jemal A. Pancreatic cancer death rates by race among US men and women, 1970-2009. J Natl Cancer Inst. 2013;105(22):1694–700. doi: 10.1093/jnci/djt292. PubMed PMID: 24203988.

14. Zhu B, Zou L, Han J, Chen W, Shen N, Zhong R, Li J, Chen X, Liu C, Shi Y, Miao X. Parity and pancreatic cancer risk: evidence from a meta-analysis of twenty epidemiologic studies. Sci Rep. 2014;4:5313. doi: 10.1038/srep05313. PubMed PMID: 24936955; PMCID: PMC4060503.

15. Lee E, Horn-Ross PL, Rull RP, Neuhausen SL, Anton-Culver H, Ursin G, Henderson KD, Bernstein L. Reproductive factors, exogenous hormones, and pancreatic cancer risk in the CTS. Am J Epidemiol. 2013;178(9):1403–13. doi: 10.1093/aje/kwt154. PubMed PMID: 24008905; PMCID: PMC3813312.

16. Andersson G, Borgquist S, Jirstrom K. Hormonal factors and pancreatic cancer risk in women: The Malmo Diet and Cancer Study. Int J Cancer. 2018;143(1):52–62. doi: 10.1002/ijc.31302. PubMed PMID: 29424426; PMCID: PMC5969235.

17. Wong A, Chan A. Survival benefit of tamoxifen therapy in adenocarcinoma of pancreas. A case-control study. Cancer. 1993;71(7):2200–3. PubMed PMID: 8384066.

18. Theve NO, Pousette A, Carlstrom K. Adenocarcinoma of the pancreas--a hormone sensitive tumor? A preliminary report on Nolvadex treatment. Clin Oncol. 1983;9(3):193–7. PubMed PMID: 6616993.

19. Satake M, Sawai H, Go VL, Satake K, Reber HA, Hines OJ, Eibl G. Estrogen receptors in pancreatic tumors. Pancreas. 2006;33(2):119–27. doi: 10.1097/01.mpa.0000226893.09194.ec. PubMed PMID: 16868476.

20. Hingorani SR, Wang L, Multani AS, Combs C, Deramaudt TB, Hruban RH, Rustgi AK, Chang S, Tuveson DA. Trp53R172H and KrasG12D cooperate to promote chromosomal instability and widely metastatic pancreatic ductal adenocarcinoma in mice. Cancer Cell. 2005;7(5):469–83. doi: 10.1016/j.ccr.2005.04.023. PubMed PMID: 15894267.

21. Rhim AD, Mirek ET, Aiello NM, Maitra A, Bailey JM, McAllister F, Reichert M, Beatty GL, Rustgi AK, Vonderheide RH, Leach SD, Stanger BZ. EMT and dissemination precede pancreatic tumor formation. Cell. 2012;148(1-2):349–61. doi: 10.1016/j.cell.2011.11.025. PubMed PMID: 22265420; PMCID: PMC3266542.

22. Li J, Byrne KT, Yan F, Yamazoe T, Chen Z, Baslan T, Richman LP, Lin JH, Sun YH, Rech AJ, Balli D, Hay CA, Sela Y, Merrell AJ, Liudahl SM, Gordon N, Norgard RJ, Yuan S, Yu S, Chao T, Ye S, Eisinger-Mathason TSK, Faryabi RB, Tobias JW, Lowe SW, Coussens LM, Wherry EJ, Vonderheide RH, Stanger BZ. Tumor Cell-Intrinsic Factors Underlie Heterogeneity of Immune Cell Infiltration and Response to Immunotherapy. Immunity. 2018. doi: 10.1016/j.immuni.2018.06.006. PubMed PMID: 29958801.

23. Casey SC, Tong L, Li Y, Do R, Walz S, Fitzgerald KN, Gouw AM, Baylot V, Gutgemann I, Eilers M, Felsher DW. MYC regulates the antitumor immune response through CD47 and PD-L1. Science. 2016;352(6282):227–31. doi: 10.1126/science.aac9935. PubMed PMID: 26966191; PMCID: PMC4940030.

24. Dennis MK, Field AS, Burai R, Ramesh C, Petrie WK, Bologa CG, Oprea TI, Yamaguchi Y, Hayashi S, Sklar LA, Hathaway HJ, Arterburn JB, Prossnitz ER. Identification of a GPER/GPR30 antagonist with improved estrogen receptor counterselectivity. The Journal of steroid biochemistry and molecular biology. 2011;127(3-5):358–66. doi: 10.1016/j.jsbmb.2011.07.002. PubMed PMID: 21782022; PMCID: 3220788.

25. Morrison AH, Byrne KT, Vonderheide RH. Immunotherapy and Prevention of Pancreatic Cancer. Trends Cancer. 2018;4(6):418–28. doi: 10.1016/j.trecan.2018.04.001. PubMed PMID: 29860986.

26. Subramanian A, Tamayo P, Mootha VK, Mukherjee S, Ebert BL, Gillette MA, Paulovich A, Pomeroy SL, Golub TR, Lander ES, Mesirov JP. Gene set enrichment analysis: a knowledge-based approach for interpreting genome-wide expression profiles. Proceedings of the National Academy of Sciences of the United States of America. 2005;102(43):15545–50. doi: 10.1073/pnas.0506580102. PubMed PMID: 16199517; PMCID: PMC1239896.

27. Wang F, Flanagan J, Su N, Wang LC, Bui S, Nielson A, Wu X, Vo HT, Ma XJ, Luo Y. RNAscope: a novel in situ RNA analysis platform for formalin-fixed, paraffin-embedded tissues. J Mol Diagn. 2012;14(1):22–9. doi: 10.1016/j.jmoldx.2011.08.002. PubMed PMID: 22166544; PMCID: PMC3338343.

